# Rapid light-dependent degradation of fluorescent dyes in formulated serum-free media

**DOI:** 10.1101/578856

**Authors:** Peter A. Morawski, Samantha J. Motley, Daniel J. Campbell

## Abstract

Chemically defined serum-free media is increasingly used as a tool to help standardize experiments by eliminating the potential variability contributed by pooled serum. These media are formulated for the culture and expansion of specific cell types, maintaining cell viability without the need for exogenous animal proteins. Formulated serum-free media could thus help improve viability and reduce variability during sample preparation for flow cytometry, yet a thorough analysis of how such media impact fluorochrome-antibody conjugates has not been performed. In this study, we expose fluorescent antibody-labeled cells or antibody capture beads to white light in the presence of various hematopoietic cell culture media and provide evidence that formulated serum-free media permit rapid light-initiated fluorescent dye degradation in a cell-independent manner. We observed fluorescence signal loss of several dyes, which included fluorescence spillover into adjacent detectors. Finally, photostability of antibody-fluorochrome conjugates in formulated serum-free media is partially restored in the presence of either serum or vitamin C, implicating reactive oxygen species in the observed signal loss. Thus, our data indicate that formulated serum-free media designed to standardize cell culture are not currently optimized for use with fluorochrome-antibody conjugates, and thus extreme caution should be exercised when using these media in cytometric experiments.

## Introduction

Cell culture using chemically defined serum-free media represents a standardized and scientifically accepted alternative to the use of pooled animal sera (1–3). Immunological studies commonly use formulated serum-free media to ensure consistent culture and expansion of hematopoietic cells, such as primary human lymphocytes and chimeric antigen receptor (CAR) T cells (4–7). Varieties of formulated serum-free media are available commercially for many immune cell types, each including defined lot-to-lot combinations of amino acids, recombinant proteins, growth factors, and inorganics. While the development and use of formulated serum-free media aims to limit technical variability in experiments, the impact of such media on antibody-conjugated fluorescent dye stability has not been well studied.

Great lengths have been taken to standardize cytometry sample preparation (8), and the efficient conjugation and proper handling of fluorescently-labeled antibodies (9). While the properties of a dye are largely unchanged following antibody conjugation (10), tandem dyes, which rely on fluorescence resonance energy transfer (FRET) between donor and acceptor fluorophores, are susceptible to oxidation, leading to less efficient FRET (11). For example, indotricarbocyanine (Cy7)-conjugated PE and APC fluorophores (12) may degrade in response to light, oxygen, or tissue fixation chemicals (11,13,14), resulting in loss of tandem dye signal intensity and introducing additional spillover into detector of the donor fluorophore. A quantitative assessment of PE tandem dyes in staining buffer revealed that increasing white light exposure resulted in proportionally elevated fluorochrome degradation, with the largest impact observed on Cy7 conjugates (11). The same study found a small but appreciable tandem dye signal loss during long-term temperature- and light-controlled storage, yet overall, shielding tandem dyes from light improved their long-term stability (13). Photon-induced fluorochrome degradation is thus a well-known issue, however an analysis of flow cytometry antibody conjugate stability in formulated serum-free media has not been performed.

In this study, we evaluated fluorescent antibody photostability in formulated serum-free media compared with traditional flow cytometry buffers. Our data demonstrate that serum-free media permit rapid light-induced degradation of fluorochromes in a cell-independent manner, while addition of serum or vitamin C limit fluorescence signal loss. Thus, although formulated serum-free media can standardize cell culture and reduce experimental variability, we find these media in their existing formulations are unreliable for use during flow cytometry due to their negative impact on fluorochrome photostability following even brief exposure to light.

## Materials and Methods

### Human tissue and cell isolation

All human samples were obtained upon written informed consent at the Virginia Mason Medical Center (Seattle, WA). All studies were approved by the Institutional Review Board of the Benaroya Research Institute (Seattle, WA). PBMC were isolated using Ficoll-Hypaque (GE-Healthcare; Chicago, IL) gradient separation. CD4^+^ T cells were enriched using CD4 microbeads (Miltenyi Biotec; Bergisch Gladbach, Germany).

### Media and reagents

The following chemically defined or partially defined serum-free media were used: X-Vivo^TM^15 Hematopoietic Cell Medium (Lonza; Bend, OR); Macrophage SFM (Thermo Fisher; Waltham, MA); AIM V^TM^ Serum-free medium (Thermo Fisher; Waltham, MA); Immunocult^TM^-XF T cell expansion medium (STEM CELL Technologies; Vancouver, Canada). Control comparisons were performed using RPMI 1640 medium with L-glutamine and phenol red (Thermo Fisher; Waltham, MA), or 1X calcium-magnesium-free PBS (Sigma Aldrich; St. Louis, MO).

### Antibodies and labeling

Human leukocytes were stained with fluorescently-labeled antibodies diluted in staining buffer made up of 1X PBS and 3% FBS for 15-20 minutes at 37°C in the dark. Antibody capture beads, UltraComp eBeads^TM^, (Thermo Fisher; Waltham, MA) were stained in the dark according to manufacturer instructions using 1X PBS. For fully-stained human samples, single color controls were generated using UltraComp eBeads^TM^ and PBMC labeled with eBioscience Fixable Viability dye eFluor780 (Thermo Fisher; Waltham, MA). The following fluorescently conjugated antibody clones were used in this study: CD4-BV421 RPA-4, CCR7-BV605 G043H7, CD103-BV605 BerACT8, CCR6-BV650 G034E3, CD4-BV650 OKT4, CD4-BV711 OKT4, CD4-BV785 OKT4, CD4-AF488 RPA-T4, CD4-FITC SK3, CCR4-PerCP/Cy5.5 L291H4, CD3-PE HIT3a, CD4-PE/Dazzle594 OKT4, CD25-PE/Cy5 BC96, CCR6-PE/Cy7 G034E3, CD8α-AF647 SK1, anti-mouse CD45.2-AF700 104, CD45RA-AF700 HI100, CD4-AF700 OKT4, CD4-APC/Cy7 OKT4, CD45RA-APC/Cy7 HI100, (BioLegend; San Diego, CA). CD45-BUV395 H130, CD28-BUV737 CD28.2, CXCR3-BV421 1C6, CD3-v450 UCHT1, CD4-BV510 SK3, CD8α-v450, RPA-T8, CD127-BV786 HIL7RM21, CD4-PE/CF594 RPA-T4, CD127-PE/Cy7 HIL7RM21, CD45-APC/H7 2D1 (BD Biosciences; San Jose, CA). CD4-PerCP/ef710 SK3, CD103-PE/Cy7 B-Ly7, CD45RA-APC/ef780 HI100 (Thermo Fisher; Waltham, MA). CCR10-PE 314305 (R&D Systems; Minneapolis, MN).

### Strategy for modulating and measuring fluorescent signal loss

After staining, fluorochrome antibody-labeled human T cells, PBMC, or antibody capture beads were resuspended in the indicated media or buffers. Fluorescent measurements of stained samples were recorded in the dark, immediately. The same samples were then exposed to either of two sources of white light: 1) the two manufacturer-installed, user-controlled, light emitting diodes (LED) contained within the sample injection chamber of a FACSAria^TM^II or FACSAria^TM^Fusion cell sorter, or 2) using ambient fluorescent light (Sylvania, 28W, 40 inch fluorescent tubes) on the laboratory benchtop (20-25°C), for a duration up to 1 hour. The fluorescence of samples exposed to the cell sorter LEDs was measured continuously in real-time during acquisition, for a duration of 3 to 7 minutes with 100 r.p.m. used for sample agitation. Where indicated, Hyclone FBS or vitamin C reagent grade L-ascorbic acid (Sigma Aldrich; St. Louis, MO) were added to the media after staining, but before exposure to light to determine their effects on fluorescent signal loss.

Fluorescent signal was assessed by measuring the change in mean fluorescence intensity over the course of sample acquisition and light exposure, and by monitoring increased spillover signal into adjacent detectors for which no staining was performed.

### Data acquisition and analysis

Data acquisition was performed as indicated using either a four-laser FACSAria^TM^II cell sorter (no ultra-violet laser), a FACSAria^TM^Fusion cell sorter or a BD^TM^LSRII analytical cytometer (BD Biosciences; San Jose, CA) each equipped with five lasers: ultra-violet (355 nm), violet (405 nm), blue (488 nm), yellow-green (561 nm) and red (637 nm). Cytometer calibration, and control fluorescent measurements were performed using Sphero^TM^ rainbow calibration particles, 8-peak (Spherotech Inc.; Lake Forest, IL). Compensation was calculated using FACS DIVA^TM^ Software (BD Biosciences; San Jose, CA) acquisition defined matrices. Data were analyzed with FlowJo 10 (TreeStar, Inc.; Ashland, OR).

### Statistical analysis

Significance was determined using an ordinary one-way ANOVA with the Dunnett posttest for multiple comparisons calculated using Prism 8.0 software (GraphPad).

## Results

### Light-dependent fluorescence signal loss in defined serum-free media can occur rapidly to select dyes in a cell-independent manner

Many published immunological studies use specially formulated serum-free media to promote standardized hematopoietic cell culture and expansion (4–7), but a thorough characterization of the interaction between these media and antibody-fluorophore conjugates has not been performed. We therefore examined how different formulated serum-free media might impact the signal stability of fluorescent antibody-labeled cells.

We first labeled freshly isolated human CD4^+^ T cells with a staining panel designed to discriminate CD4^+^ naïve and memory T_helper_ subsets. Fluorescently labeled CD4^+^ T cells were resuspended in formulated serum-free hematopoietic media, X-Vivo15^TM^, and their fluorescence intensity was recorded immediately in the dark (Fig. 1A, t=0m). These cells were then exposed to the LED within the sample injection chamber of the cell sorter for a minimum of five minutes and fluorescence intensity was recorded again (Fig. 1A, t=5m). All analyses were performed on live cells, indicated by an absence of signal for fixable amine-reactive dyes (data not shown). We observed an approximate 10-fold loss of staining intensity for CD45RA-AlexaFluor700. CD127-PE/Cy7 signal loss was similar and included a reciprocal increase of signal into the detector for the PE donor fluorophore, which greatly impacted T_helper_ subset discrimination. Other dyes, such as the polymers BV605 and BV650, remained intact following exposure to the LED. These data suggest that formulated serum-free media reduces photostability of select fluorochrome-antibody conjugates, even in response to abbreviated light exposure. We next asked whether serum-free hematopoietic media formulated for use with different immune cell populations also impact dye photostability. For this, CCR6-PE/Cy7 and CD4-AF488 dual-labeled PBMC were resuspended in either staining buffer made of PBS-FBS, one of four distinct formulated serum-free media, or serum-free RPMI (RPMI-sf). PE/Cy7 and AF488 fluorescence intensity, as well as CCR6-PE/Cy7 spillover signal into the PE detector (Fig. 1B, t=0h) were measured immediately in the dark using an analytical cytometer. These cells were not stained with PE conjugates. Labeled cells were then exposed to ambient light on the benchtop for 1 hour at room temperature. Measurements were then repeated for PE/Cy7 and spillover into its donor fluorophore detector, PE (Fig. 1B, t=1h). When cells were kept in staining buffer during 1 hour of fluorescent light exposure, signal intensity remained constant for CCR6-PE/Cy7 and CD4-AF488, with no spillover signal increase in the PE detector. In contrast, PE/Cy7 photostability was reduced by varying degrees depending on whether antibody-labeled cells were kept in either serum-free X-VivoTM15, Macrophage_SFM_, AIM V^TM^, ImmunoCult^TM^-XF, or RPMI-sf (Fig. 1B, 1C). In some serum-free media, for example AIM V^TM^, tandem dye signal spillover into the adjacent PE detector was extensive despite a limited change in CCR6-PE/Cy7^+^ signal (Figs. 1B, 1C), indicating that increased donor fluorophore signal could occur in the absence of much noticeable loss of FRET. These data demonstrate that formulated serum-free media permitted light-induced fluorochrome-antibody signal loss to a subset of tested dyes, including substantial spillover into adjacent detectors.

**Figure 1.**
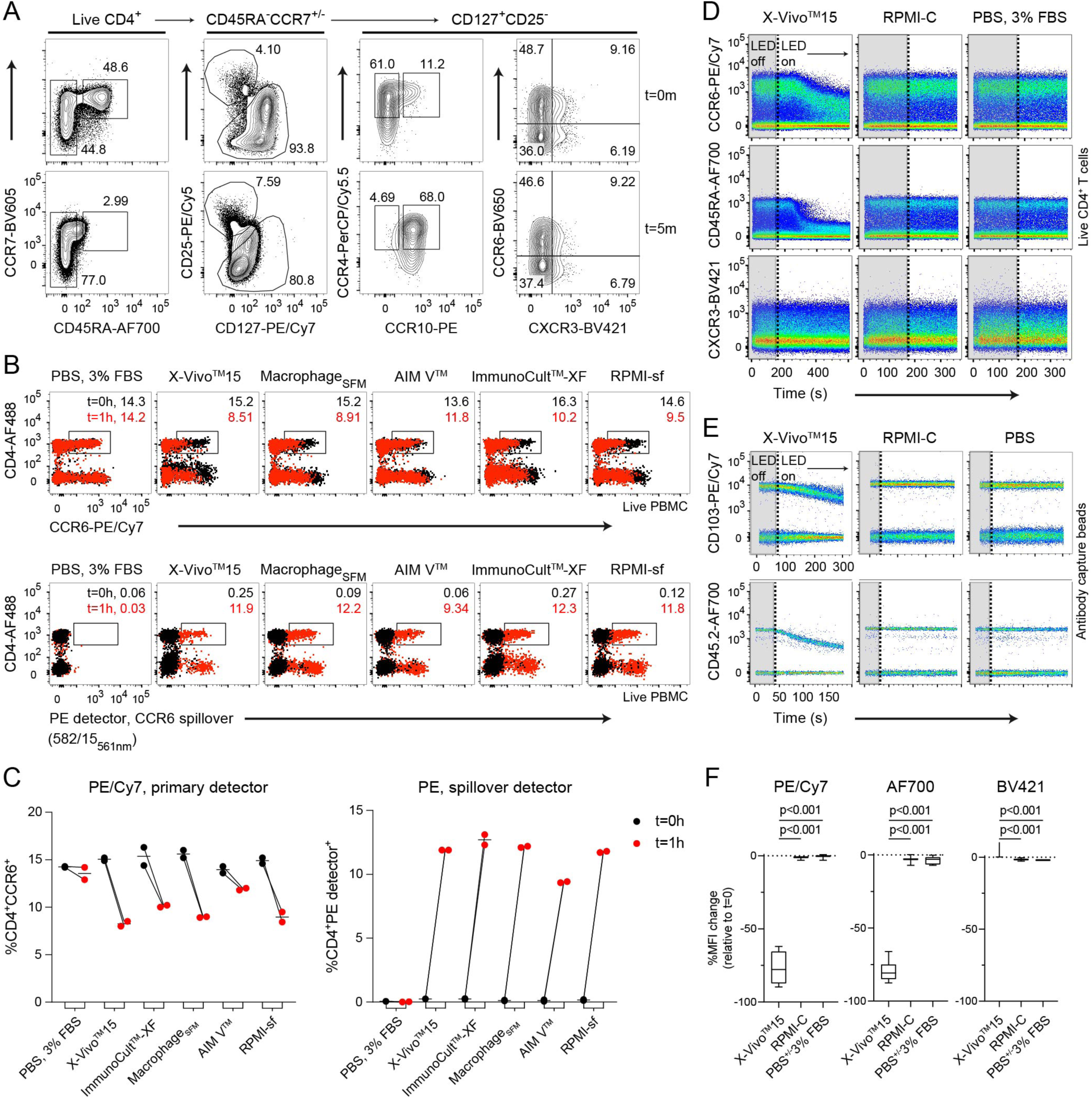
Rapid light-initiated, cell-independent fluorochrome degradation in formulated serum-free media. (**A**), Fluorescence measurements of labeled CD4^+^ T cells resuspended in formulated serum-free medium, X-Vivo^TM^15, either before (t=0m) or after exposure to the LEDs (t=5m) in the cell sorter sample injection chamber. Cells gated as indicated aside each plot. (**B**, **C**), Fluorescence measurements of labeled, live gated PBMC resuspended in the indicated media were taken before (t=0h, black dots) or after (t=1h, red dots) exposure to ambient fluorescent light on the benchtop. The gated frequency for each timepoint is shown. Simultaneous measurements were performed for (*top*) CCR6-PE/Cy7 into its primary detector and (*bottom*) spillover into the PE detector. The bandpass filter and laser line of the spillover detector are indicated. (C) A graphical summary of the data showing changes to CCR6-PE/Cy7 signal and concomitant spillover into the PE detector across both timepoints. (**D-F**), Real-time fluorochrome photostability of either single-labeled (D) CD4^+^ T cells or (E) antibody capture beads was compared across the indicated media or buffer. Surface protein expression was measured first briefly in the dark (“LED off”, gray boxes ending in vertical dotted line), and then in response to light (“LED on”). (F) A statistical comparison of percent MFI change following up to five minutes of light exposure is shown for pooled cells and antibody capture beads. Data are representative of at least 2 independent experiments. Significance was determined by ordinary one-way ANOVA with Dunnett’s posttest for multiple comparisons.

We next wanted to assess the rate of light-initiated fluorescence loss in formulated serum-free media. Fluorescence measurements of stained human T cells were recorded in real-time on a cell sorter, first for three minutes in the dark, followed by an additional three to seven minutes of LED exposure in the sample injection chamber. We observed no signal loss on labeled T cells during the dark period, however drastic reductions to signal of PE/Cy7- and AF700-conjugates were detected within 30 seconds of LED exposure in formulated serum-free X-Vivo^TM^15 media (Fig. 1D). By comparison, BV421-conjugates were unaffected by light.

Tandem dye degradation can be impacted by metabolic processes and cell viability (14). We thus wanted to test whether the fluorescence signal loss we observed (Figs. 1A-1C) was cell-dependent. To do this, we measured the fluorescence stability of single-stained antibody capture beads in real-time for up to one minute in the dark, followed by three to five minutes of LED exposure in the cell sorter as described above. CD103-PE/Cy7 and CD45.2-AF700 labeled beads remained photostable in the dark, but a rapid signal reduction of both conjugates was observed after LED exposure (Fig. 1E). While PE/Cy7 and AF700 conjugates were susceptible to light-mediated degradation in formulated serum-free medium compared to photostable BV421 conjugates, all dyes were photostable in complete RPMI containing serum or in PBS, which was evident using either stained cells or antibody capture beads (Fig. 1F). To investigate photostability over the full visible spectrum, we single-labeled antibody capture beads with 25 different fluorescent antibody conjugates, resuspended them in X-Vivo^TM^15 serum-free media, and tested dye photostability in real-time (Fig. 2A). We observed varying degrees of light-induced signal loss to several tandem dyes including CCR6-PE/Cy7, as well as conjugates of APC/Cy7, BV785, and BV786, while base fluorophores such as PE, APC, and BV421, were largely photostable. In addition, tandem dyes previously reported to be susceptible to light-mediated degradation in other systems such as PE/Cy5 (11), were unaffected in our experimental conditions. Unexpectedly, conjugates of AF700, which is not a tandem dye, were highly susceptible to light-induced signal loss in formulated serum-free media, as indicated by decreasing MFI signal over time (Fig. 2B). These data collectively demonstrate that photostability of fluorescent antibody-labeled samples is reduced in formulated serum-free media. This phenomenon is cell-independent and impacts a select range of both tandem and base fluorophores.

**Figure 2.**
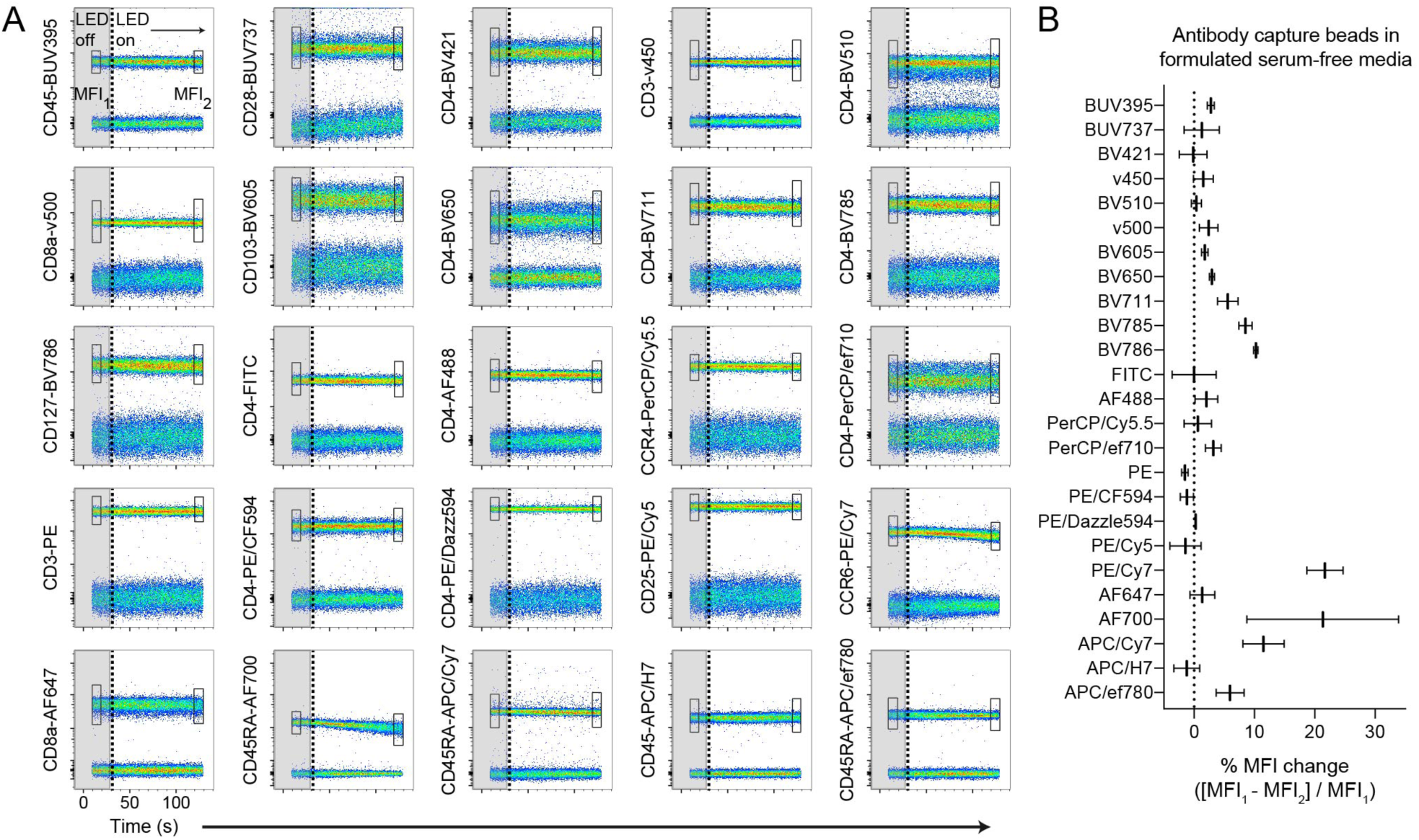
Broadly assessing fluorochrome photostability in formulated serum-free media. (**A**), Antibody capture beads single-stained with each of the 25 indicated antibody-dye conjugates were resuspended in X-VivoTM15 serum-free media. Real-time fluorochrome photostability was measured, first briefly in the dark (“LED off”, gray boxes ending in vertical dotted line), and then in response to light (“LED on”). The MFI of the positively-stained bead population was calculated for the beginning (box “MFI_1_”) and end (box "MFI_2_”) of the time course. (**B**), Graph of percent MFI change calculated from (A), following 2-3 minutes of light exposure. Data are representative of at least 2 independent experiments.

### Serum and vitamin C can limit light-induced fluorochrome degradation in formulated serum-free media

Our data indicate that fluorescent antibody conjugates are photostable in complete RPMI containing serum (Figs. 1D-1F), while a previously published study demonstrated that vitamin C could prevent APC tandem dye signal loss on stained cells (14). To test the impact of serum and vitamin C on cell-independent fluorescence signal loss, we generated single-stained antibody capture beads using PE/Cy7, APC/Cy7, or AF700 conjugates. Stained beads were placed in formulated serum-free media alone, or in the presence of either 1mM vitamin C or 3% FBS, and exposed to ambient light on the benchtop for 1 hour at room temperature. Following light exposure, we observed fluorescence signal loss of all tested antibody conjugates, and substantial spillover into the base fluorophore detectors for PE/Cy7, APC/Cy7 dyes when in formulated serum-free medium alone (Figs. 3A, 3B). Signal loss and concomitant spillover was reduced in the presence of vitamin C or FBS, though the degree to which these media additives increased photostability varied among dyes. We were unable to detect AF700 spillover into any other fluorescence detectors despite marked signal loss in the primary detector (Fig. 3B).

**Figure 3.**
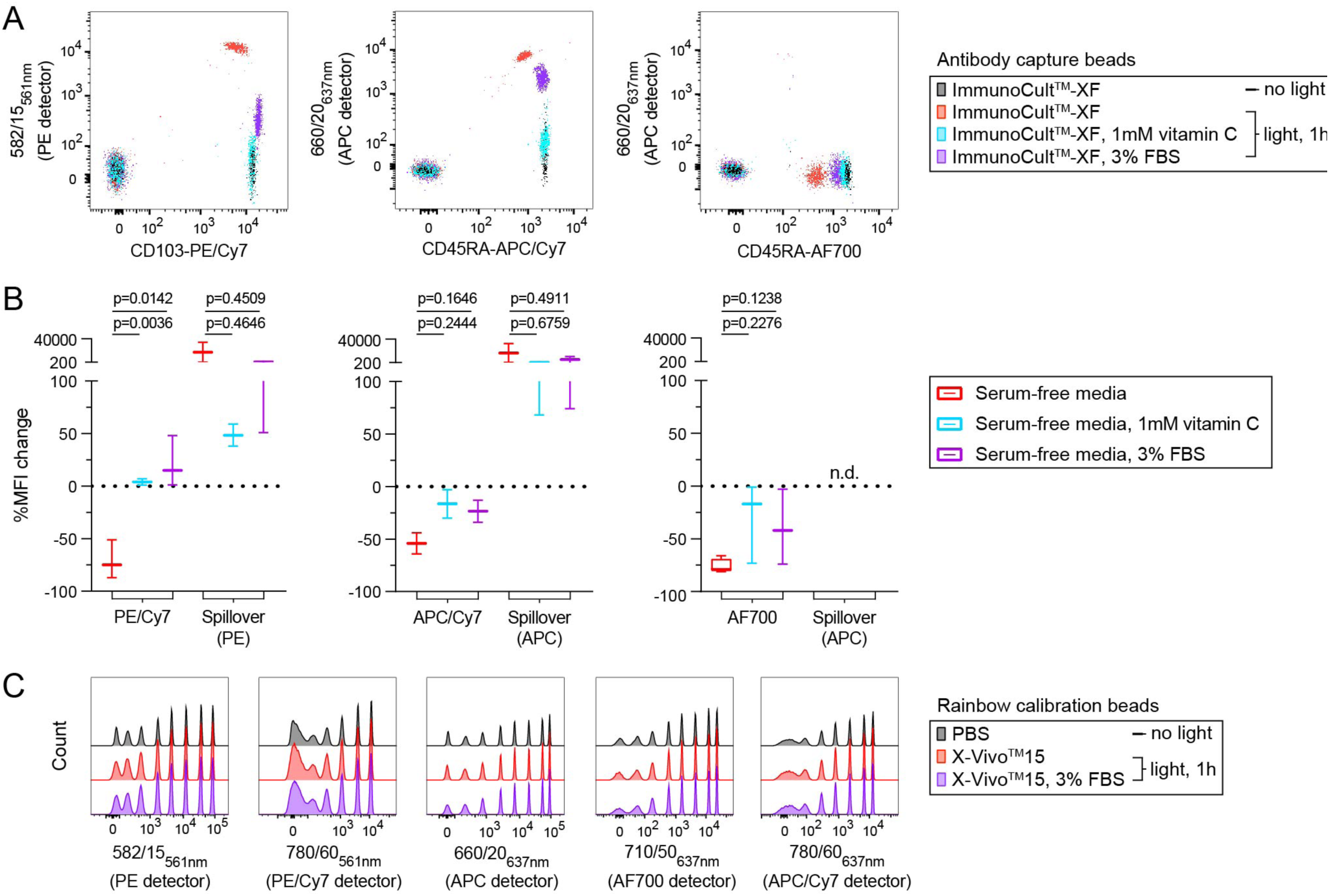
Serum and vitamin C limit light-initiated fluorochrome degradation in formulated serum-free media. (**A**, **B**), Antibody capture beads single-stained with the indicated conjugates were resuspended in formulated serum-free media, either ImmunoCult^TM^-XF or X-Vivo^TM^15. (A) Fluorescence signal and spillover into adjacent detectors was measured immediately (“no light), and again following exposure to ambient fluorescent light (“light, 1h”) with or without either vitamin C or FBS. (B) A statistical comparison of percent MFI change following light exposure is shown for data pooled from stained cells and antibody capture beads resuspended in either ImmunoCult^TM^-XF or X-Vivo^TM^15 serum-free media. (**C**), Rainbow calibration 8-peak beads were placed in PBS or formulated serum-free media, X-Vivo^TM^15 with or without FBS. Fluorescence signal was measured before and after light exposure as in (A) and for the same five detectors. Detector bandpass filter and laser line are shown. Significance was determined by ordinary one-way ANOVA with Dunnett’s posttest for multiple comparisons. Data are representative of at least 2 independent experiments. (n.d., not detectable)

To rule out that the observed fluorescence signal changes were not artifacts of formulated serum-free media, antibody lot, or the light source used, and to confirm normal function of our cytometers, we measured the fluorescence signal of 8-peak rainbow calibration beads resuspended in PBS or serum-free media. Because the calibration beads include highly stable fluorescent particles contained within the beads themselves, they should be immune to light-dependent fluorescence decay. Consistent with this notion, rainbow calibration particles were unaffected by exposure to 1 hour of ambient fluorescent light, irrespective of the buffer or media used, or the presence of serum (Fig. 3C).

## Discussion

Chemically defined serum-free media help standardize cell culture, circumventing lot-to-lot variability inherent to pooled human or animal serum. Cell type-specific formulations of serum-free media are now commonly used to permit immune cell culture without sacrificing cell viability or function. For example, these media have become a critical tool for the culture and expansion of CAR-T cells (6,7). Cytometry core facilities and technical manuals on cell sorting suggest using buffered, serum-supplemented media to help maintain cell viability during the sorting and collection of live stained cells (15), but few, if any, manufacturer guidelines or publications address the use of formulated serum-free media for these procedures. Our study thus aimed to characterize the stability of fluorescent antibodies in formulated serum-free media.

Fluorescent antibody conjugates, are susceptible to signal loss from light exposure or reactive oxygen species (ROS) (11,14). In this study, we found that white light caused rapid signal loss in serum-free media to Cy7 and select BV tandem dyes, as well as the non-tandem, AF700 (Fig. 1). Other tandem conjugates such as Cy5, which are subject to photostability issues, were not impacted under our tested conditions. Our data suggest this fluorescence signal loss is cell-independent, as it occurs not only to stained cells, (Figs. 1A-1D) but also to labeled antibody capture beads (Figs. 1E, 2). Importantly, signal loss occurred even after brief periods of light exposure that might be encountered during normal sample preparation and handling.

Not all tandem dyes behaved similarly when exposed to light in formulated serum-free media. For example, antibody capture beads labeled with the polymer tandems BrilliantViolet^TM^ 605 and 650 were more photostable in serum-free media when compared to BV785/6 (Fig. 2). The reason for this variable dye stability remains unclear, but could be connected to differences in their specific chemistry. Surprisingly, we found rapid and substantial light-induced signal loss of multiple AF700 conjugates despite the known photostability of the Alexa Fluor family of dyes (16). No other Alexa Fluor dyes we tested exhibited this effect. It remains unclear why AF700, which is not a tandem dye, was so strongly impacted by light in formulated serum-free media. Our understanding of how sample handling can impact fluorochrome degradation will be aided by the advent of spectral cytometers, which can measure the full fluorescence profile of dyes instead of only instrument-specific bandpass filters.

Two recent studies showed that cell sorting can induce oxidative stress, altering the basal metabolic state of cells (17), and that the presence of photo-sensitive molecules within standard serum supplemented cell culture medium could drive morphological changes or cell death via oxidative stress (18). In each case, addition of serum or antioxidants reduced measurable cell stress. We observed that using AIM V^TM^ media there was only a marginal loss of PE/Cy7 FRET efficiency compared with other serum-free formulations, though extensive spillover into the PE detector remained. AIM V^TM^ contains human serum albumin, a protein component similar to serum, with noted antioxidant capacity (*new ref. 19*), which could explain the small photostability increase we observed in this media. However, as the commercially manufactured media we used contain proprietary preparations it is unclear if other formulations tested contain human serum albumin, and thus difficult to resolve the noted difference in signal loss.

Vitamin C can also reduce oxidative stress by scavenging free oxygen radicals, and is sufficient to prevent cell-dependent APC tandem dye degradation (14). We find that RPMI containing serum maintains the photostability of stained cells (Figs. 1D-1F), and supplementing formulated serum-free media with serum or vitamin C can reduce, but does not eliminate signal loss and spillover into adjacent detectors (Figs. 3A, 3B). Though we did not test for the presence of ROS in our experiments, the effect of vitamin C we observe, and its published role as a scavenger of these compounds (14) suggests a light-dependent mechanism through which components of formulated serum-free media are transformed to promote fluorochrome degradation. Though we did not test cell viability after photodegradation occurred, we demonstrate that signal loss in this study is cell-independent, suggesting that any deleterious role for ROS persists independently of effects on cell viability. Further study is required to validate and dissect these mechanisms, and if necessary, address the matter of light-induced oxidative stress in serum-free media. However, care must be taken in the selection and addition of new components to sorting media. For example, vitamin C has been shown to interact with components of cell culture to produce detrimental byproducts (20). Other chemicals can also protect against oxidative stress, including selenium and 2-ME (1), though in our hands, addition of 2-ME to X-Vivo^TM^15 did not prevent light-induced fluorescence signal loss (data not shown). Additional work is required to identify specific additives that can safely and effectively address the issue of light-dependent fluorophore degradation in formulated serum-free media.

Collectively, our findings suggest that chemically defined and formulated serum-free media in their current manufactured forms do not sufficiently support the photostability of fluorochrome-antibody conjugates. We conclude that these media should be avoided during cytometric experiments, as even supplementing with serum did not fully maintain dye photostability. Even sorting cells into formulated serum-free media could prove problematic, as some sorters are equipped with lights near the collection tubes that cannot be turned off by the user, and could thus impact the reliability of post-sort purity checks. Future modification of formulated serum-free media to support dye photostability would help reduce variability of flow cytometry sample preparation and collection by eliminating the need for pooled serum and other undefined additives that could impact downstream readouts.

## Acknowledgements

We thank Jessica Hamerman, Sabine Spath and Liliane Khoryati for helpful discussion and review of the manuscript.

## Author Contributions

P.A.M. and D.J.C. designed the experiments. P.A.M. developed the methods and wrote the manuscript. P.A.M. and S.M. performed experiments. P.A.M., S.M., and D.J.C. reviewed and edited the manuscript. D.J.C. supervised the project.

